# A module-based approach for post-omics, post-GWAS network-based gene classification

**DOI:** 10.1101/2025.08.11.669721

**Authors:** Alexander McKim, Christopher A. Mancuso, Arjun Krishnan

## Abstract

Complex traits and diseases are highly polygenic and understanding the full set of genes involved is a central challenge in biomedicine. However, due to sample size limitations and noise (technical and biological), experimental approaches for disease-gene discovery such as transcriptomics and GWAS result in long, noisy, heterogeneous gene lists, which may be trimmed to a subset of likely relevant genes while leaving several false negatives. Computational gene classification approaches, especially those using genome-scale molecular interaction networks, are promising avenues for complementing such experimental findings by analytically expanding observed gene lists based on the functional relatedness between genes. We previously introduced the network-based gene classification approach, *GenePlexus*, which was rigorously benchmarked to show state-of-the-art performance, especially for predicting novel genes associated with biological processes and fine-grained phenotypes. Network-based gene classification performance, however, declines for diseases, especially when the inputs are omics and GWAS-based long gene lists. Here, we show that such disease gene lists span multiple biological processes spread across the molecular network and propose *ModGenePlexus*, a new network-based gene classification method that takes a two-stage approach. First, clustering and semi-supervised learning decomposes the input gene list into coherent denoised network gene modules. Then, *ModGenePlexus* trains supervised (*GenePlexus*) classifiers for each module and aggregates predictions to return genome-wide rankings. We benchmarked *ModGenePlexus* across simulated data, transcriptomic signatures, and GWAS datasets (together spanning hundreds of diseases), showing improved recovery of known disease genes compared to *GenePlexus*. Beyond improved classification, the results of enrichment analysis of *ModGenePlexus* outputs are much more interpretable by virtue of revealing nuanced biological processes. Together, these results establish *ModGenePlexus* as a scalable, interpretable tool for gene classification of GWAS and -omics derived genelists across diverse biological contexts.

## Introduction

Determining the genes that play a role in a given cellular pathway, fine-grained (developmental, anatomical, behavioral) phenotype, or complex trait/disease is a grand challenge in the fields of genetics and molecular biology. In the case of pathways and phenotypes, careful experimental discovery is still the gold standard for addressing this challenge, but it is labor intensive and does not scalable across the genome. Therefore, computational gene classification methods have emerged as critical alternatives that allow researchers to more readily prioritize pathway/phenotype gene annotations^1–8^ for experimental follow-up. These methods aim to expand known gene sets by identifying novel genes likely to share functional roles, helping address the question: *what other genes are involved in my pathway/phenotype of interest?* Because a gene’s role is highly dependent on its functional context, ‘network-based’ gene classification methods that leverage the functional relationships (i.e., molecular network connections) among genes to predict novel pathway/phenotype annotations have proven widely successful^9,10^. Our group previously developed *GenePlexus*^1,11^, a network-based approach that builds a supervised machine learning (ML) model trained on the network relationships of known genes and uses this model to classify novel genes, with each gene associated with a probability based on the similarity of its global network connectivity to that of the known gene list. Based on extensive benchmarking across diverse gene sets of cellular pathways, diseases, and traits, we showed that this supervised learning approach outperforms the classical label propagation method that spreads information from known genes along the network edges to infer novel genes^12^. *GenePlexus* performs well when applied to smaller, coherent (highly inter-connected) gene sets such as biological processes and fine-grained phenotypes^13,14^. However, the performance of network-based gene classification declines when the inputs are larger, noisy, heterogeneous gene lists generated by the most widely-used approaches for revealing disease-associated genes: transcriptomic profiling and GWAS. Each transcriptomics study or GWAS typically produces hundreds to thousands of genes associated with the complex trait or disease of interest but such large gene lists are often substantially noisy, containing false positives and false negatives, making it difficult to identify truly disease-relevant genes without resorting to applying an extremely stringent statistical threshold. A central reason for this noise is the inherent low statistical power of omics studies, which stems from a fundamental imbalance: the number of samples (*n*) is small relative to the vast number of features or hypotheses being tested (*p*), such as tens of thousands of genes in 3–12 samples (transcriptomics) or millions of SNPs in few hundred thousands individuals (GWAS). This *n vs p* disparity limits the resolution of discovery, leading to incomplete and error-prone gene lists. Specifically, In transcriptomics, statistical power is highly dependent on the number of replicates^15^, but most RNA-seq experiments have an insufficient number of replicates^16^. In GWAS, increases in cohort size over time have consistently yielded new discoveries, and yet, it is well acknowledged in the field that GWAS results contain false positives and false negatives^17,18^ because of low power. These challenges and observed limitations in omics based studies are explained and compounded by the extreme polygenicity of complex traits. For example, complex traits and disease are rarely driven by a single pathway; instead, they reflect the perturbation of multiple, distinct molecular processes that emerge as the disease. The vast multi-dimensional nature of many biological traits and diseases, and the difficulty for omics studies to give powered results are why gene classification and predicting new genes for even large experimental gene lists is critical for uncovering the full biological landscape of mechanisms behind complex traits. *GenePlexus* is ideal for prioritizing relevant genes and pathways in a network-driven manner because biological networks capture how disease-associated genes work together to orchestrate pathways/processes relevant to disease phenotypes, and it relies on identifying a single coherent network pattern connecting the input genes. However, large gene lists that span multiple distinct pathways involved in the disease are often distributed across different regions of the genome-wide interaction network, making it unlikely that a single, unified network pattern can fully capture the underlying disease biology. *GenePlexus’s* performance thus suffers due to the assumption that all input genes are functionally related, the presence of irrelevant genes and absence of relevant ones, and the lack of mechanism for deconvolving functional heterogeneity embedded in the input.

To address these challenges, we propose *ModGenePlexus*, a new network-based gene classification framework that recognizes that omics-based large gene lists associated with complex traits and diseases often encompass multiple distinct biological processes^19^. Therefore, instead of trying to find one consistent pattern in the underlying gene network for the entire trait/disease gene list, *ModGenePlexus* takes a divide-and-conquer approach by taking advantage of the fact that, in a gene network, biological processes and pathways are captured as local network neighborhoods^20^. *ModGenePlexus* refines large input gene lists by clustering them into smaller, biologically coherent modules and applying network-based classification separately to each module. By focusing on local network neighborhoods, *ModGenePlexus* improves accuracy by denoising the input and recovering novel genes linked to specific disease-relevant processes.

We validated *ModGenePlexus* through a series of systematic evaluations using a study-bias holdout validation scheme (see ***Methods***) on large, simulated and real-world datasets to demonstrate its robustness in diverse gene classification scenarios. First, we conducted simulations using “traits” constructed by combining multiple biologically meaningful gene sets to assess performance for varying gene set sizes and noise levels. These simulations confirm that *ModGenePlexus* outperforms *GenePlexus*, particularly in settings with heterogeneity or false positive genes. We then applied *ModGenePlexus* to differentially-expressed genes from hundreds of public transcriptomic datasets^21^ and demonstrated substantial performance improvements over *GenePlexus* across a wide range of diseases, drug treatments, and gene perturbations. Next, we evaluated *ModGenePlexus* on GWAS data, demonstrating that it accurately predicts significantly associated genes, indicating its ability to navigate the inherent noise and redundancy in GWAS-derived gene lists. Finally, to illustrate the interpretability and added value of *ModGenePlexus*, we present a case study on type 2 diabetes, which demonstrates how its inherent modular approach provides insight into distinct gene clusters, uncovering processes missed by non-modular (single-model) classification applied to traits with highly complex genetic architecture. Together these results highlight the advantages of a module-based approach for gene classification with large, noisy, heterogeneous gene lists from high-throughput experiments and emphasize the broader implications of network-based modularity for interpreting and generating novel hypotheses from complex biological datasets generated by large-scale -omics studies.

## Methods

### Data processing

#### Network processing

To cluster and to generate network-based features for the supervised learning model we leveraged the widely-utilized STRING^22^ function network throughout this work. We downloaded v10 of the homo sapiens network (*9606.protein.links.detailed.v12.0.txt)* directly from the STRING website and retained only edges with an edge weight > 0.7. The Ensembl protein IDs were mapped to Entrez IDs using mygene.info^23^, and in the case where one Ensembl protein ID mapped to more than one Entrez ID we added an edge for each Entrez ID. This resulted in a network with 16,624 nodes and 400,729 edges. The STRING network includes information from a variety of different resources (physical interactions, coexpression, gene annotation databases, etc) and is best suited to handle the complex functions and phenotypes seen in the large gene sets in our evaluation set.

#### Processing gene set collections

We evaluated *ModGenePlexus* using a diverse set of gene sets, incorporating Gene Ontology Biological Processes in our simulations and gene sets from CRowd Extracted Expression of Differential Signatures^21^ (CREEDS) and GWAS Atlas^24^ for real-world validation. A common processing step for all gene set collections involves removing genes from any gene set if it is not found in the processed STRING network. For more in-depth information on the gene set collections see **File S1**.

#### Gene ontology biological processes

The Gene Ontology Biological Process^25,26^ (GOBP) gene set collection was compiled from utilizing gene annotations from Mygene.info^23,27,28^. These direct annotations were then propagated using the hierarchical structure of the GO ontology. Genes were filtered based on their presence in STRING to align with network-based analyses. Additionally, GOBP terms were refined by retaining only those containing a minimum of 10 genes and a maximum of 100 genes. To simulate complex traits composed of multiple distinct biological processes, we randomly combined 2, 5, and 10 GO term gene annotations to create simulated traits. False positive rates of 0.0, 0.25, 0.50, 0.75, and 1.0 were applied, where the rate represents the percentage of the size of the simulated trait. Genes that were not annotated to any GO term in the simulated trait were added as false positives to evaluate *ModGenePlexus* under varying noise levels.

#### CREEDS database

We compiled gene sets from the CRowd Extracted Expression of Differential Signatures (CREEDS) database (v1.0), which includes manually curated genes from GEO^29^. We included only manually curated signatures and retained gene sets with at least 100 genes, ensuring sets are sufficiently complex. We used three distinct data modalities: single gene perturbations (𝑛 = 972), single drug perturbations (𝑛 = 234), and diseases (𝑛 = 311).

#### GWAS Atlas database

We compiled gene sets from the GWAS Atlas (v20191115) utilizing MAGMA^30^ gene prioritization scores for three different thresholds; 𝑝 < 1×10^−2^ (𝑛 = 670), 𝑝 < 1×10^−5^ (𝑛 = 32), and 𝑝 < 1×10^−8^ (𝑛 = 15), only retaining sets with at least 100 genes. We chose to create GSCs for multiple p-value thresholds from MAGMA because while it is common to use very strict thresholds for GWAS data due to loss of significance from multiple hypothesis correction and the assumption that these top genes capture all the underlying biology of a trait, it has been shown that even genes determined to be less significant can have biological meaning, especially when accounting for their cumulative effects^31^.

### Inner-working of *ModGenePlexus*

#### Gene set clustering

We used DOMINO^32^ to generate the modules used by *ModGenePlexus* because it is one of the best methods for finding “active modules” ^33^, *i.e.*, subnetworks within a large genome-scale network that capture active molecular processes specific to a condition of interest, probed by the analyzed transcriptomics or GWA study^34^. DOMINO uses Louvain clustering^35^ to partition the genome-wide network by optimizing modularity. DOMINO then filters the initial partitions based on a list of genes and applies label propagation to refine the clusters, making them more topologically coherent. This process removes some genes from the initial set through filtering while incorporating additional genes via label propagation.

### Validation Scheme

To evaluate model performance in a way that reflects the real-world challenge of novel gene discovery, we use a study bias holdout scheme. This evaluation is intentionally stringent, as it tests the ability to predict understudied genes, which tend to have fewer or weaker network connections compared to well-characterized genes^36,37^. For all gene sets, the initial positive genes are those that are originally annotated in the gene set. The selection process then differs between methods:

- *GenePlexus*: All originally annotated genes are positive labels
- *ModGenePlexus*: Genes contained in a given cluster after running DOMINO.

For each input gene set (trait), we assess gene study bias by counting the number of PubMed abstracts^38^ each gene appears in. Genes in the top two-thirds of PubMed mention frequency were designated well-studied and assigned to the positive training set. The bottom one-third, representing understudied genes, are held out as the positive test set. All models use this same positive training/test split to ensure consistency across evaluations. Negative genes were identified using the approach implemented in PyGenePlexus^11^. This method defines negative examples as genes not annotated to any gene set in a reference collection that significantly overlaps (p < 5×10⁻², one-sided Fisher’s exact test) with the input gene set. This process also identifies a set of neutral genes that are not used in training or evaluation. Negative genes were then randomly divided into training and test sets, maintaining the same training-to-test ratio used for positive genes. Importantly, the negative test set consists only of genes that are consistently identified as negative across all models, including both *GenePlexus* and *ModGenePlexus*, ensuring that no negative test gene is used during training in either *GenePlexus* or *ModGenePlexus*.

#### Running the supervised learning model

GenePlexus and *ModGenePlexus* were run using a modified version of PyGenePlexus (v1.0.1)^11^ Python package. All models were trained using the STRING network with adjacency matrix-based feature representations. PyGenePlexus uses a logistic regression with L2 regularization implemented in scikit-learn for classification.

#### Aggregating module gene predictions

For *ModGenePlexus*, gene scores across modules were aggregated across all identified modules using the tau score^39^, defined by formula:

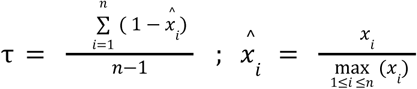

Where τ is the tau score, 𝑥*_i_* is a gene’s rank for module 𝑖, and 𝑛 is the total number of genes for which *ModGenePlexus* returns predictions. Each gene’s final score was computed by aggregating its predictions across all identified modules within a given gene set. Genes were then ranked based on these scores, and the performance of test genes in this ranked list was compared to the baseline performance of *GenePlexus*. τ was chosen as the metric to define genes based if they have noticeably high prediction scores in a module, implying the gene is relevant in functional processes.

#### Evaluation metric

Evaluations were conducted using log2(auPRC/prior), which is the area under the Precision-Recall Curve (auPRC) normalized by the prior probability of a gene having a positive label. auPRC and prior are defined as follows:

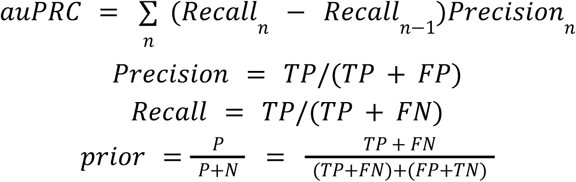

Where TP is true positives, FN is false negatives, FP is false positives, and TN is true negatives. P and N are the ground truth labels. This approach accounts for class imbalance, ensuring model comparison remains robust despite differences in positive and negative label distributions^40^.

#### GOBP enrichment of gene sets

GOBP enrichment was performed using the clusterProfiler^41^ R package. Enrichment was conducted separately for *GenePlexus* and for each individual module model prediction in *ModGenePlexus*. For module-based predictions, genes with a probability of >= 0.80 were considered predicted positives. For *GenePlexus*, due to the larger input sizes, enrichment was conducted using either a 0.80 probability threshold or a threshold that limited the number of predicted genes to at most twice the original input size, whichever resulted in fewer genes. All enrichment analyses were conducted using the org.Hs.eg.db (3.20.0) annotation package, with default parameters. Significance was determined using a q-value threshold of 0.05.

#### Normalized beta coefficients of *(Mod)GenePlexus* models

To compare the contribution of the most important features across models, we extracted the raw logistic regression coefficients from the trained *GenePlexus* and *ModGenePlexus* classifiers and created heatmaps using the ComplexHeatmap and the circlize^42,43^ R packages. To enable direct comparison across models and modules, we applied mean-centered scaling followed by max-absolute-value scaling, normalizing the coefficients within each model to a common range of [-1,1]. Specifically, for each model, each coefficient was centered by subtracting the mean across all coefficients, and then divided by the maximum absolute centered value to facilitate comparison across modules:

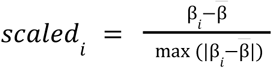

Where β*_i_* is the beta coefficient the 𝑖^th^ beta coefficient in the list of model coefficients, and β is the mean beta coefficient value. We visualize the top 20 scaled model coefficients across each module and *GenePlexus*, where each coefficient in this case applies to the top 20 gene predictions for each model.

#### Calculating information content of GOBPs

To assess the specificity of enriched GOBPs, we calculated the information content (IC)^44^ for all enriched terms:

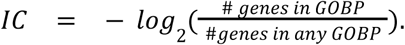

## Results

### Single-model gene classification leads to poor performance for heterogeneous input gene lists

Following our previous work, we re-evaluated *GenePlexus* using diverse data sources, including biological process-gene, disease-gene, and trait-gene associations. While *GenePlexus* performs well on curated datasets, its performance significantly declines when applied to experimental datasets such as transcriptomics-derived gene lists (**Figure 1A-B**). These gene lists present unique challenges due to noisy signals, where some relevant genes may not exhibit differential expression (false negatives), while some observed changes may not reflect causal or functionally relevant biology (false positives), making robust predictions more difficult. The correlation between edge density and performance reflects this issue (Figure 1B): gene sets derived from biological processes tend to have high edge densities and are often localized within a local network neighborhood, while gene sets from GWAS exhibit lower edge densities and are more scattered in the network. Further, despite increased edge density in transcriptomic-derived datasets, performance does not improve relative to other modalities, highlighting the limitations of *GenePlexus*’s single-model approach for large and highly complex gene sets. For network-based methods, the localization of genes within the network is a key predictor of performance. In STRING, biological processes (**Figure 1C**) tend to be more localized than disease- or GWAS/transcriptomics-derived datasets (**Figure 1D-E**). These limitations and performance drops are not unique to *GenePlexus*; similar patterns of performance decline are observed in other network-based gene classification methods, including the classical label propagation approaches^1^.

**Figure 1:**
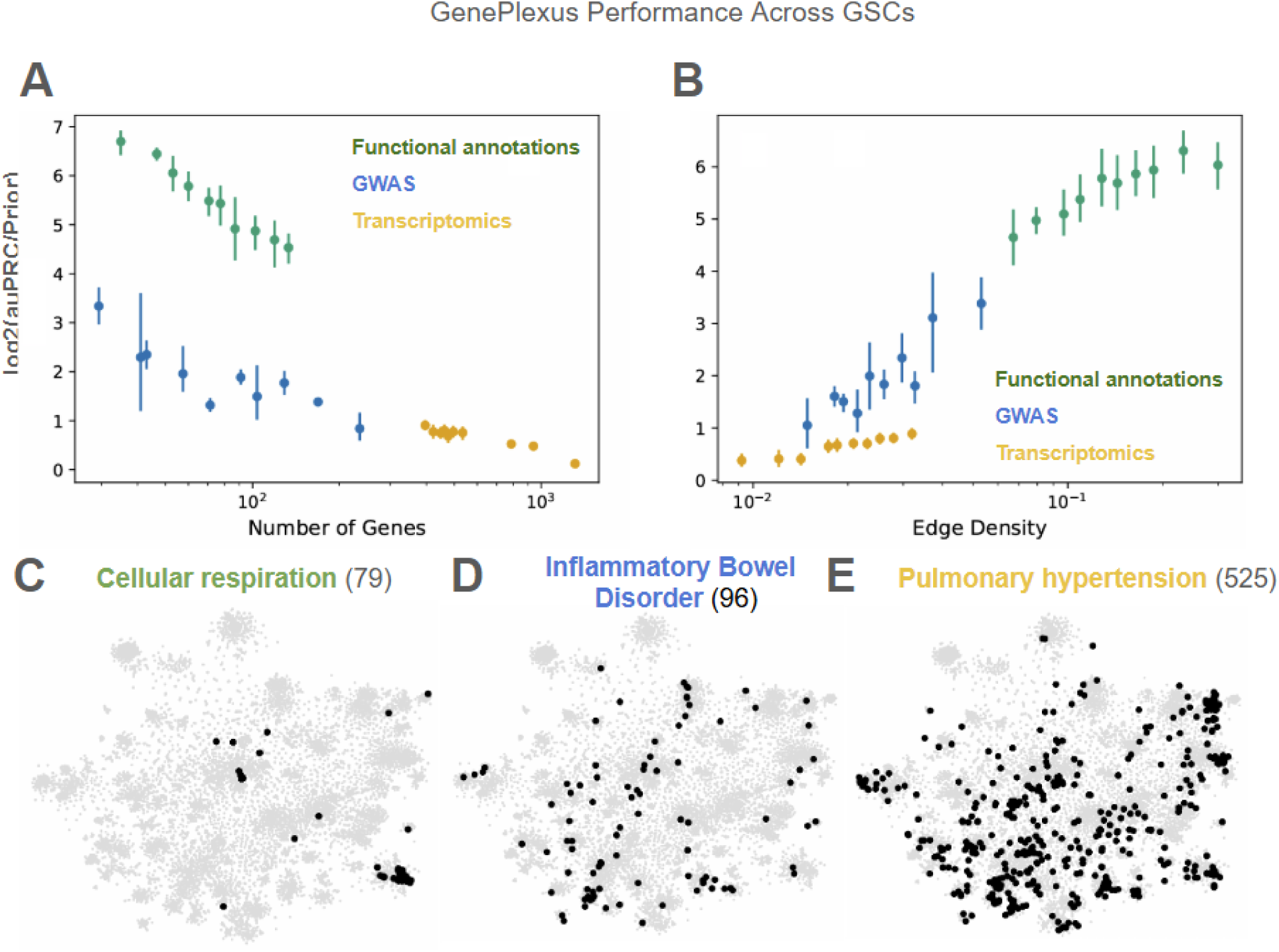
GenePlexus performs poorly on large, dispersed gene sets. (A) GenePlexus performance as a function of gene set size across different data modalities. Larger gene sets, particularly from transcriptomic datasets, exhibit consistently poor performance. (B) GenePlexus performance as a function of network edge density across data modalities. Despite increased edge density in transcriptomic-derived gene lists, performance does not improve relative to other modalities, highlighting its limitations in large, complex gene sets. (C) The localization of genes of an average size biological process. (D) The genes of an average sized GWAS set. The genes are not as localized within the network. (E) The genes of an average sized transcriptomics set.

### *ModGenePlexus*: a network-based divide-and-conquer ML approach to tackle long, noisy, heterogeneous gene lists

To address the limitations of single-model gene classification, we developed *ModGenePlexus,* a modular extension of *GenePlexus* designed to improve performance for large, noisy, and heterogeneous gene lists derived from omics experiments (**Figure 2**).

**Figure 2:**
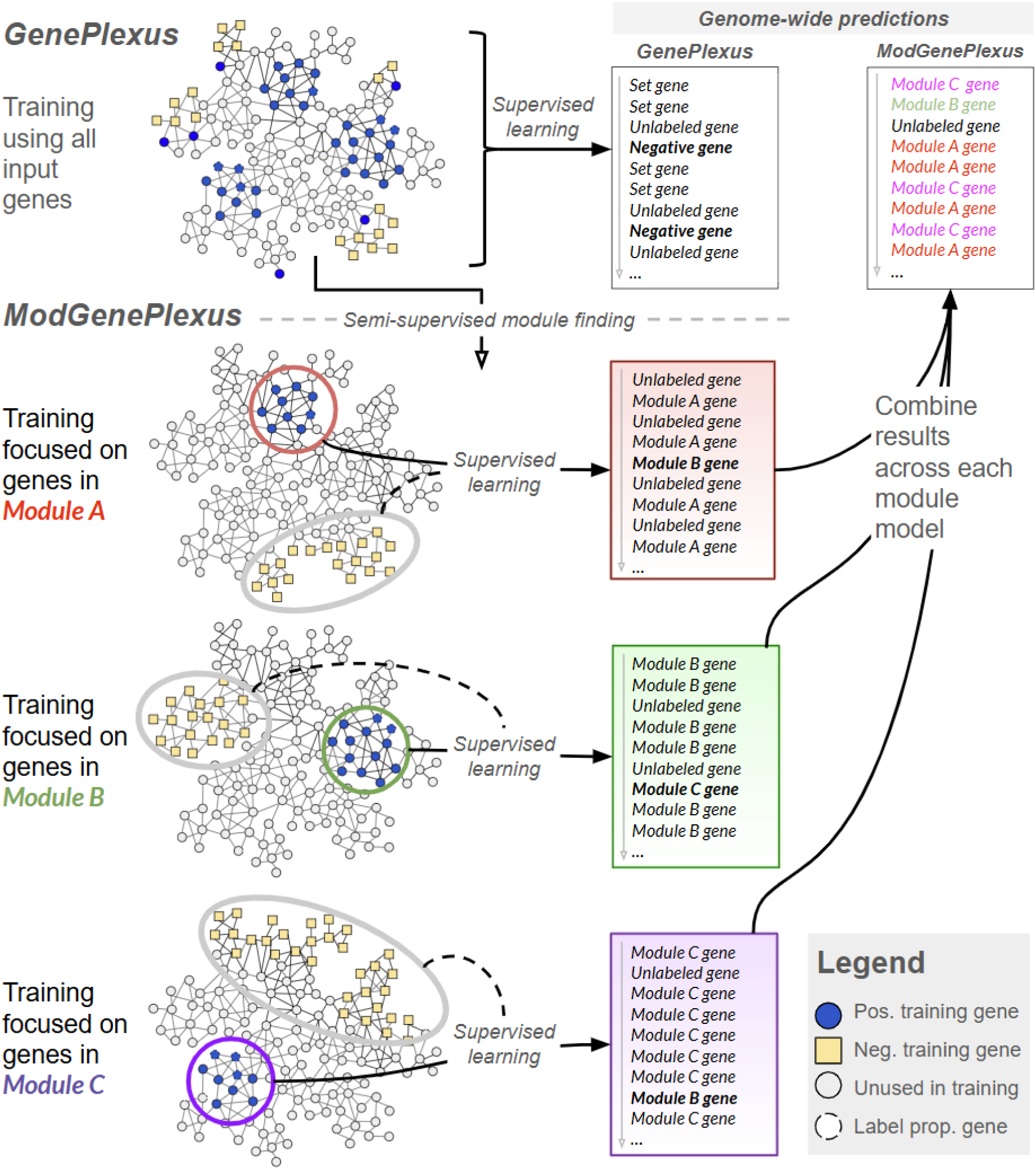
The ModGenePlexus method. Genes are clustered to discover modules, where semi-supervised learning filters out genes that do not belong to sufficiently large modules. Genes outside the original input set but located in disease-enriched modules are incorporated as additional inputs. These refined cluster gene sets are then processed by GenePlexus, which builds a logistic regression model for each module to classify genes at the module level. The individual module predictions are subsequently aggregated to generate a final gene prediction score.

*ModGenePlexus* works by first clustering the long list of trait/disease-associated genes into smaller, denoised, topological and biologically coherent subsets that we refer to henceforth as ‘modules’. Semi-supervised label propagation is then applied to identify and expand genes well-connected to each discovered module. Finally, network-based gene classification, *i.e.*, *GenePlexus*, is applied independently to each expanded module, finally presenting the top genes for each module and an aggregated top gene prediction across modules. *ModGenePlexus* improves gene classification accuracy by refining the input gene list by removing weakly connected genes and expanding it to include new candidates with strong module-specific connectivity. This dual refinement is particularly useful for gene lists derived from low-powered and/or minimally preprocessed datasets where false positives and false negatives may be prevalent. We systematically validated *ModGenePlexus* across simulated, transcriptomics-derived, and GWAS-derived gene lists, demonstrating substantial improvements in gene classification performance compared to *GenePlexus*. Additionally, by identifying distinct gene modules, *ModGenePlexus* enhances biological interpretation and reveals context-specific pathways underlying complex traits and diseases.

### *ModGenePlexus* accurately recovers genes associated with simulated heterogeneous traits

One challenge in designing and validating *ModGenePlexus* was the lack of a gold-standard disease module set^45^. To address this challenge, we constructed large gene sets (simulated “traits”) composed of biologically meaningful “modules” using sets of genes annotated to GOBP terms. Each trait consisted of either two, five, or ten GOBPs, treating each GOBP as an individual gene module. To mimic false positives and real-world noise, we introduced randomly sampled non-GOBP genes, where false positives were introduced where the number of false positives is 0.25, 0.50, and 1.0 times the number of genes in the original set. *ModGenePlexus* consistently outperformed *GenePlexus* where performance was measured using log2(auPRC/prior) (see ***Methods***; **Figures 3, S1**).

**Figure 3:**
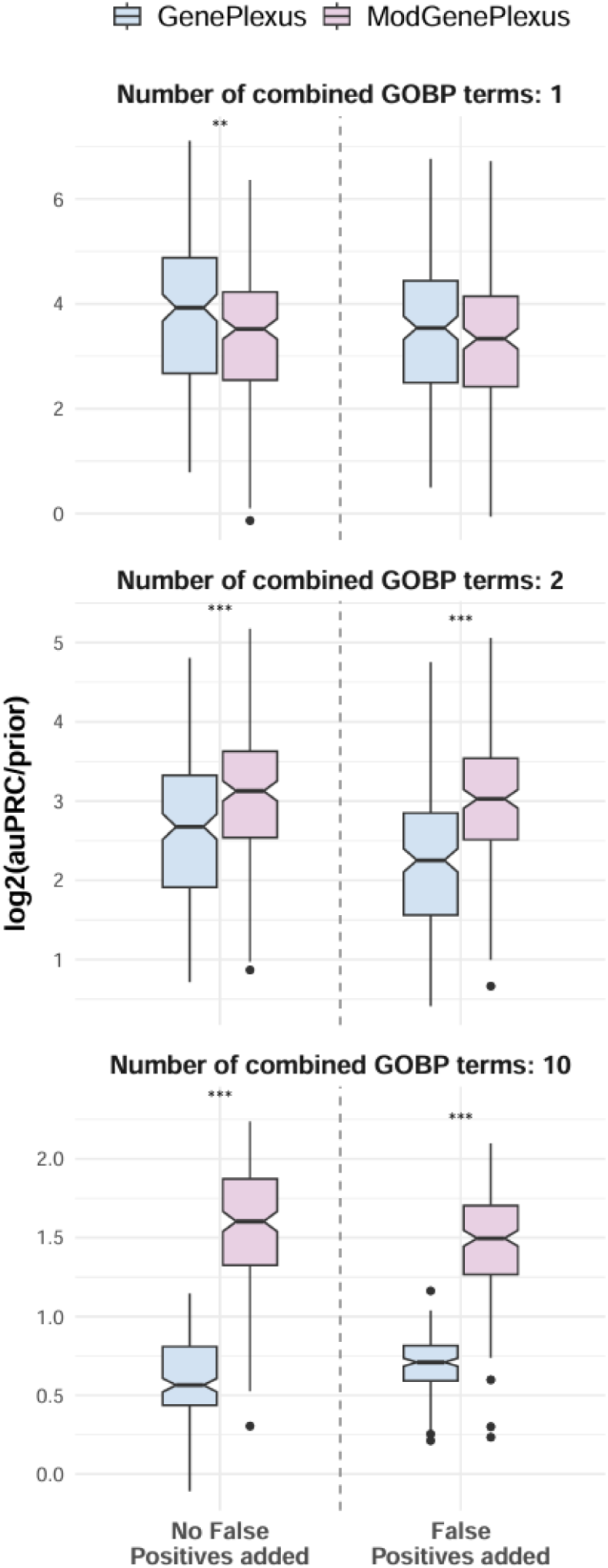
ModGenePlexus outperforms GenePlexus on simulated traits generated by combining multiple GOBP terms. Models were tested on individual GOBP terms and simulated traits composed, 5 or 10 combined terms. False positives were introduced where the number of false positives added is 50% of the simulated set size Results show that as the number of combined GOBP terms increases, the performance gap between ModGenePlexus and GenePlexus widens, with ModGenePlexus consistently outperforming GenePlexus, particularly for multi-term traits.

As the number of combined GOBP terms increased and false positives grew, *ModGenePlexus*’s advantage over *GenePlexus* became more pronounced. The only exception was when testing a trait made of a singular GOBP – in which case *GenePlexus* performed better when no false positives were present, but both methods performed similarly as false positives were introduced. This outcome aligns with *GenePlexus*’s established ability in classifying genes for individual GOBPs. Overall, *ModGenePlexus* excels when analyzing datasets with multiple distinct biological processes or when false positives are prevalent.

### *ModGenePlexus* outperforms the single-model approach in gene classification of transcriptomics-derived gene lists

We designed *ModGenePlexus* to handle real-world gene-based results from omics-scale experiments, the most common being large, noisy lists of differentially expressed genes (DEGs) from RNA-seq or microarray experiments. To test our method’s effectiveness in this setting, we evaluated *ModGenePlexus* on the DEG results of a manually-curated collection of public microarray experiments studying three types of ‘conditions’: diseases, gene perturbations, and drug treatments. These DEG gene lists pose an ideal challenge for *ModGenePlexus* because they often contain hundreds or thousands of gene associations and high levels of noise^46,47^. After filtering based on size and experiment count, we obtained 311 disease gene lists, 972 gene perturbation gene lists, and 234 drug treatment gene lists with ≥100 genes total and ≥10 understudied genes in the positive test genes (see ***Methods***). We first tested the full *ModGenePlexus* pipeline, which integrates module finding, denoising, semi-supervised gene expansion, and supervised gene classification (via *GenePlexus*).

Again, using *GenePlexus* as our baseline, across all types of conditions, *ModGenePlexus* significantly outperforms *GenePlexus* (**Figure 4A**). This better performance holds even when we skip the semi-supervised gene expansion step, showing the importance of roughly breaking down large, complex gene lists into modules (**Figure S2**). We additionally calculated the edge density of the original gene set and the obtained modules. The average edge density of the modules is most often greater than the edge density of the original gene set used for *GenePlexus* (**Figure 4B**), which confirms that the modules are more cohesive in the underlying network. We also tested the importance of the denoising step by pooling all genes contained in any module into a single gene list and then using that combined gene list to train a single *GenePlexus* model (**Figure S3**). We see that while pooling the denoised modules into a single gene list leads to some improvement to *GenePlexus*, it’s performance still falls way short of *ModGenePlexus*, reinforcing the advantage of analyzing individual modules separately rather than treating the gene list that contain multi-factorial biology as a single unit. Lastly we analyzed the performance of predicted DOMINO genes performance and comparing to *ModGenePlexus* results. DOMINO fails to recover most positive genes and *ModGenePlexus* demonstrated superior classification accuracy (**Figures S4-S5**).

**Figure 4:**
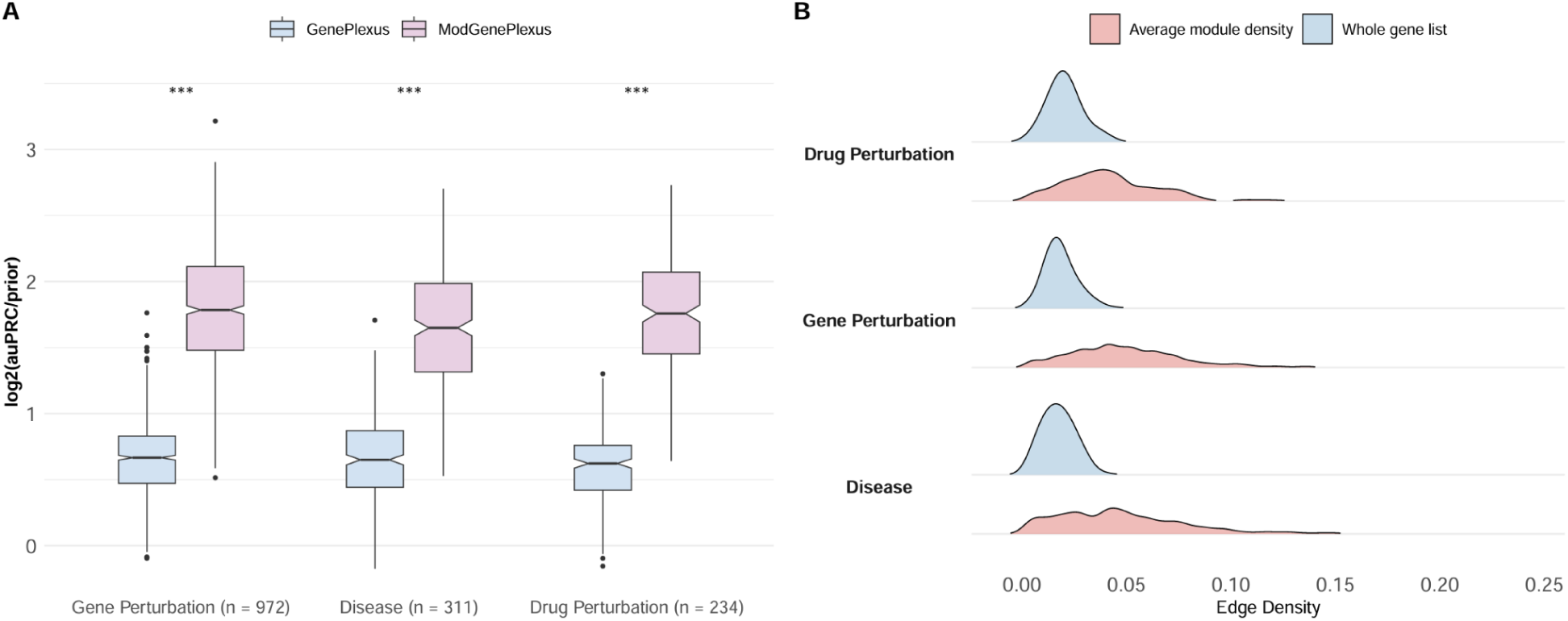
ModGenePlexus outperforms GenePlexus for transcriptomic gene lists. (A) ModGenePlexus significantly outperforms GenePlexus across three transcriptomic gene set collections from CREEDS: gene perturbations, disease, and drug perturbations. Stars refer to a Wilcoxon rank-sum test, where significance values are p=6.49e-295, p=1.63e-87, p=5.08e-75 respectively. (B) Distribution of network edge densities for GenePlexus input (whole gene set) and the average edge density for modules (Average module density). Module edge densities are typically higher than the edge density of the whole initial set.

### ModGenePlexus outperforms the single-model approach in gene classification of GWAS-derived gene lists

The second major application area for *ModGenePlexus* is to reprioritize and interpret trait- or disease-associated genes from a GWAS. Compared to analyzing the gene-based results of functional omics studies like transcriptomics, the application to GWAS presents unique challenges due to stringent multiple hypothesis corrections of summary statistics at the level of SNPs and genes (derived using algorithms like MAGMA^30^). These challenges intensify as GWAS sample sizes increase because more significant hits are constantly discovered^48^. To assess the performance of *ModGenePlexus* on GWAS results, we applied it to gene-based summaries for a wide range of traits and diseases generated by MAGMA, compiled in GWAS Atlas^24^. We used three significance thresholds (liberal, nominal, and stringent) — 𝑝 < 1×10^−2^, 𝑝 < 1×10^−5^, and 𝑝 < 1×10^−8^ to derive gene lists associated with each trait/disease and retained gene lists with both ≥100 genes and had statistically significant modules returned from DOMINO, which resulted in 645, 31, and 15 traits/diseases for each threshold, respectively.

For all three thresholds, *ModGenePlexus* outperformed *GenePlexus* (**Figure 5A**). Notably, *GenePlexus* performed considerably worse on GWAS data with the most stringent threshold, often yielding performance worse than random. In an alternative validation approach, we trained models using genes defined at the liberal and nominal thresholds (𝑝 < 1×10^−2^and 𝑝 < 1×10^−5^) but evaluated them on understudied genes defined at the nominal threshold. These two thresholds were selected because they yielded enough sufficiently-sized gene lists to support robust model training and evaluation. The 𝑝 < 1×10^−5^ *ModGenePlexus* model performed best (**Figure 5B**), with the 𝑝 < 1×10^−2^ model also surpassing *GenePlexus* when evaluated at 𝑝 < 1×10^−5^. These findings show that *ModGenePlexus* leverages module discovery to filter out poor-quality hits more effectively than using a strict p-value threshold to compile a gene list. By reducing the impact of false positives while retaining biologically relevant genes, *ModGenePlexus* enhances GWAS-based gene classification, particularly for understudied and less significant genes.

**Figure 5:**
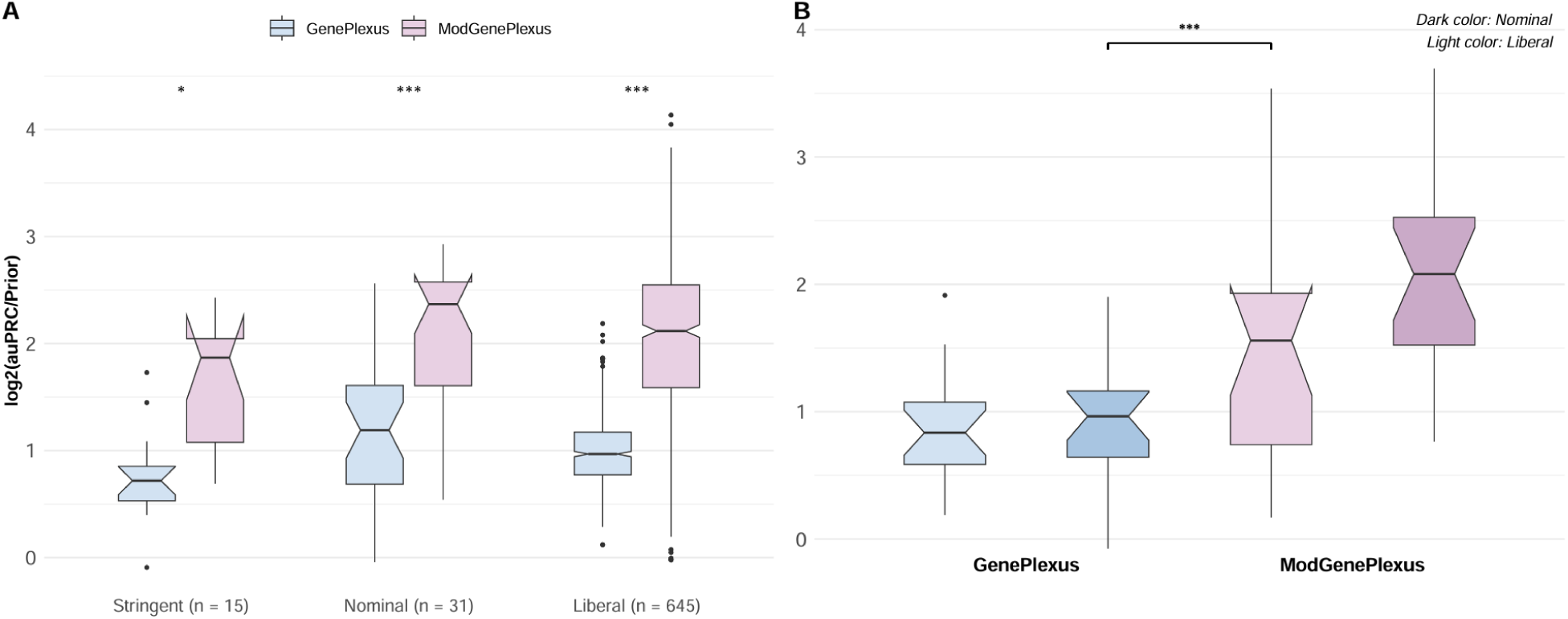
ModGenePlexus outperforms GenePlexus on GWAS gene level predictions. (A) Performance comparison using MAGMA-derived gene scores from GWAS summary statistics at three significance thresholds (Stringent: p < 1e-8, Nominal: p < 1e-5, and Liberal: p < 1e-2). ModGenePlexus consistently outperforms GenePlexus across all thresholds, though both methods exhibit a decline in performance at the strictest threshold (Stringent). Stars refer to a Wilcoxon rank-sum test, where significance values are p=1.32e-4, p=8.93e-7, p=1.01e-131 respectively (B) ModGenePlexus models trained on less stringent GWAS thresholds (Nominal, Liberal) demonstrate improved classification performance when evaluated on 1e-5 test genes, outperforming the standard GenePlexus model. Wilcoxon sum rank test p < 0.001.

### *ModGenePlexus* intrinsically learns local network neighborhoods and provides substantially interpretable enrichment results

To confirm *ModGenePlexus*’s ability to isolate specific signals within large gene lists, we examined the model’s learned coefficients which offers insight into the network-based signals driving its predictions. We visualized the beta coefficients from the supervised learning models created in *GenePlexus* and *ModGenePlexus* to determine if beta coefficient values were more extreme or more specific in *ModGenePlexus* models compared to *GenePlexus*. *ModGenePlexus* highlighted smaller, high-confidence gene subsets which suggests context-specific biological relevance. In contrast, *GenePlexus* distributed lower-weighted coefficients across a broader gene list (**Figure 6A**).

**Figure 6:**
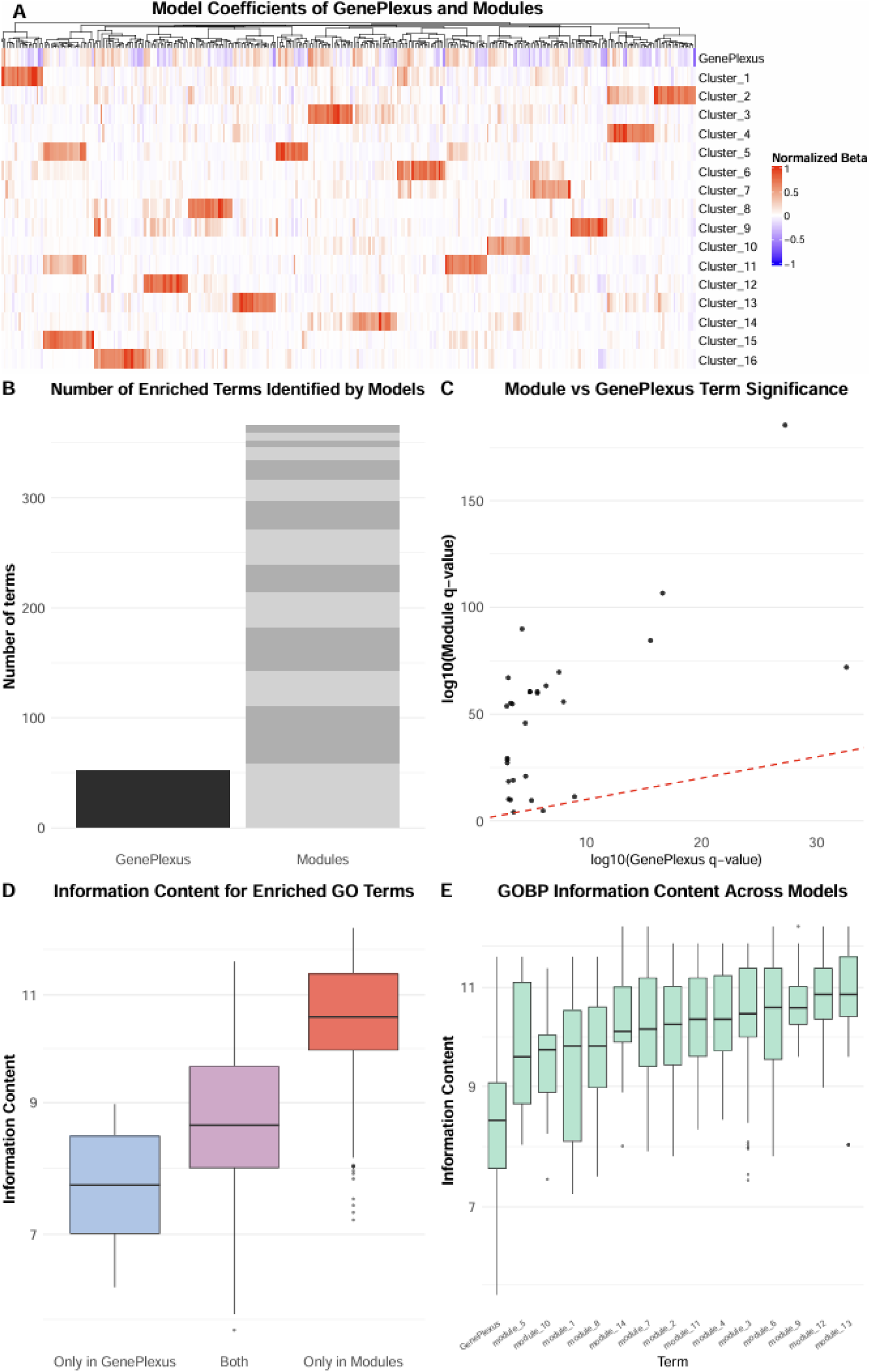
Using modules uncovers relevant biology that a full gene set misses. (A) Normalized logistic regression model coefficients (min-max scaled) for GenePlexus and ModGenePlexus. ModGenePlexus modules exhibit more concentrated weight values within trait-specific modules, whereas GenePlexus coefficients are more diffuse and less significant. (B) Number of enriched GOBP terms identified by GenePlexus and ModGenePlexus. The module-based approach results in substantially more enriched terms compared to GenePlexus. (C) Enrichment values of GOBP terms shared between GenePlexus and ModGenePlexus. Module-level enrichment values in ModGenePlexus are significantly higher than those observed in GenePlexus predictions based on the entire gene list. (D) Information content distribution of enriched GOBP terms categorized by enrichment in only GenePlexus, only ModGenePlexus, or both. Terms enriched in both methods, as well as those enriched only in GenePlexus, have smaller information content than those enriched exclusively in ModGenePlexus, reflecting its finer biological resolution. (E) Comparison of enriched GOBP information content between GenePlexus and ModGenePlexus modules. Module-based enriched terms have higher information content than those identified using the entire gene set.

The primary approach for interpreting long gene lists from high-throughput studies is to perform gene set enrichment analysis to identify overrepresented pathways/processes, which may be relevant to the condition being studied. Therefore, we decided to assess the impact of gene classification and reprioritization by *ModGenePlexus* on enrichment-based biological interpretability. To do so, we compared the results of GO Biological Process (GOBP) enrichment on the top-ranked genes from *ModGenePlexus* and *GenePlexus*, using Type 2 diabetes as a case study. This disease has a highly heterogeneous signature, with ∼2000 genes across 14 CREEDS-derived differential expression studies. *ModGenePlexus* improved gene classification performance, prompting further analysis of downstream enrichment. Multiple correction was performed (Benjamini-Hochberg; *p < .001*) was used to determine significance. *ModGenePlexus* leads to recovering significantly more enriched terms than *GenePlexus*, including many captured only by *ModGenePlexus* (**Figures 6B**). Furthermore, terms shared between both methods show stronger enrichment under *ModGenePlexus* (**Figure 6C**), suggesting higher specificity and precision. For example, GOBP cellular response to insulin was discovered by *ModGenePlexus* (*BH=4.10e-06)* but not *GenePlexus* (*BH=4.63e-01)* To test whether *ModGenePlexus* yields more focused biological insights, we compared the size of enriched gene lists. Module-level enrichment from *ModGenePlexus* produced more compact, functionally distinct terms (**Figure 6D-E**), while *GenePlexus*-enriched terms had less information content. Terms unique to *GenePlexus* also had less information content than those shared between methods (**Figure 6C**). Shared terms also tended to have higher information content than those enriched exclusively in *ModGenePlexus* results. In total, *ModGenePlexus* yielded 366 unique enriched terms compared to 52 from *GenePlexus* (**Figure S6**), and some terms such as oxidative phosphorylation were enriched across multiple modules (**Table S1**) suggesting that modules with distinct genes work together in the same or similar pathways. Together, these observations indicate that *ModGenePlexus* enhances interpretability by emphasizing high-resolution, modular signals rather than high-level associations seen in typical enrichment analysis of gene-based outputs from high-throughput experiments. To explore these differences further, we mapped the top-ranked predictions from both *GenePlexus* and *ModGenePlexus* to discover relevant and diverse processes by *ModGenePlexus* but not *GenePlexus*, displayed in enrichment tables (**Tables S1-S6**).

## Discussion

Improved prioritization of trait/disease-associated genes from large-scale experimental datasets using genome-scale gene networks presents significant challenges due to gene set size, noise, and biological complexity. Standard approaches, including label propagation (semi-supervised learning) and *GenePlexus* (supervised learning), that treat the entire gene list as a single unit struggle to classify genes in such cases^1^, necessitating new methods. Here, we present *ModGenePlexus*, a novel approach that addresses key limitations in network-based supervised learning with large, noisy, heterogeneous gene lists by leveraging modular structure present in gene networks, filtering out poorly connected genes (likely false positives), and producing not one but multiple reprioritized gene lists, each corresponding to network-defined biologically coherent biological processes. While several existing gene reprioritization methods^49^ generate a single genome-wide ranking of genes for a trait or disease, this single output can be difficult to interpret in practice. Given the high polygenicity of complex traits/diseases, important disease-associated genes may appear far below the top of such rankings—well beyond the intuitive cutoffs researchers typically consider for follow-up (e.g., top 10, 50, or even 100 genes). Furthermore, these single-list outputs still require manual interpretation of individual genes or post-hoc grouping based on pathway enrichment. Because in *ModGenePlexus* each gene list is explicitly linked to a network module and the corresponding enriched pathways, interpreting these results for follow-up studies becomes more straightforward. Thus, *ModGenePlexus*’s modular outputs are able to substantially elevate the interpretability and practical utility of the gene prioritization results.

We demonstrated that *GenePlexus* performs poorly on large experimental gene sets, particularly those from transcriptomic studies, where high noise and poor network localization weaken classification accuracy of single-model methods. Because *GenePlexus* applies supervised learning on the full original gene set, it cannot function as a standalone solution for these datasets. This limitation underscores a fundamental limitation of network-based classifiers: their reliance on globally coherent patterns is incompatible with the heterogeneous biological reality of polygenic traits. We showed that *ModGenePlexus*’s advantage grows as gene sets increase in size and contain more false positives. As the number of distinct perturbed pathways and processes increases, *ModGenePlexus* effectively separates distinct biological modules and improves gene classification in more complex traits. *ModGenePlexus* significantly improves gene classification for results from transcriptomic datasets. DEG lists from such datasets are not only large but also highly noisy, lacking strong network localization. By integrating filtering, semi-supervised propagation, and supervised learning, *ModGenePlexus’s* ability to filter noise and propagate meaningful gene associations makes the input gene set more topologically and biologically coherent and enables more biologically meaningful predictions for real omics data.

We use DOMINO as the method to cluster the large input gene list into modules. In this step, DOMINO also “denoises” the gene list using an approach akin to semi-supervised learning, dropping genes that are not strongly connected and including new genes in the network that are critical for the connectivity of the remaining genes. We demonstrate that using this semi-supervised learning step improves *ModGenePlexus* results and that this improvement occurs by just using the module-based breakdown of the original genes even without the newly-included genes (**Figure S2**). Therefore, though we used DOMINO in this study, in principle, any network clustering method that partitions a large input gene list into optimal modules^50^ can be used in *ModGenePlexus*.

Similar to its performance on gene lists derived from transcriptomics studies, *ModGenePlexus* enhances gene reprioritization for GWAS datasets. We show that training the model with genes selected based on nominal p-value thresholds improves the performance compared to training on genes from liberal thresholds. On the other hand, training on genes from the strictest threshold (p < 1×10^-8^) resulted in comparatively lower performance. This performance decrease likely results from the small number of retained genes that represent disparate pathways and processes involved in the trait/disease (i.e., one or a couple of genes from most of the relevant pathways rather than multiple genes from any one pathway), meaning they are unlikely to be close together in the underlying network. When the threshold is relaxed to a nominal value, other genes in relevant pathways start entering the initial list and coalesce around the top genes in the underlying network, forming more cohesive modules that *ModGenePlexus* can leverage. Previous studies demonstrate that genes with more nominal p-values can still have biological relevance^31^, meaning strict thresholds may remove false positives but also introduce false negatives. Solely relying on strict genome-wide significance thresholds may systematically underrepresent functionally important genes. *ModGenePlexus’s* network-based refinement offers a principled solution to this longstanding challenge in GWAS interpretation.

Beyond superior performance in reprioritizing genes, using Type 2 Diabetes as a case study, we show how *ModGenePlexus* can lead to greater biological insights compared to *GenePlexus*. *ModGenePlexus* uncovers more granular biological pathways (**Figure 6**) by leveraging modular network structure and the ability to use module-specific model results, rather than the aggregation into a single list. For example, from the list of genes differentially expressed in Type 2 Diabetes, *ModGenePlexus* uniquely identified enrichment in copper ion transport^51–54^ in module 5 (**Table S2**) and iron homeostasis pathways^55–58^ in modules 1 and 3, which the single-model approach failed to detect (**Tables S3-5**). Distinct insulin-related^59–61^ pathways (enriched in modules 7 and 12) were only detected by *ModGenePlexus* (**Table S5**). These findings illustrate how *ModGenePlexus* reveals biological processes that would be overlooked by considering all disease genes as a single unit.

## Conclusion

*ModGenePlexus* provides a scalable and biologically meaningful approach to prioritize trait/disease-associated genes from large, noisy, heterogeneous gene lists derived from high-throughput experiments. By leveraging modular network structure and integrating filtering, propagation, and supervised learning, *ModGenePlexus*’s divide-and-conquer approach significantly outperforms single-model methods like *GenePlexus* in network-based prioritization of genes from transcriptomic and GWAS datasets. Ultimately, its ability to refine gene associations and uncover biologically meaningful pathways makes *ModGenePlexus* a powerful tool for post-omics hypothesis generation across diverse disease studies.

## Author contributions

A.K., A.M., and C.A.M. conceived and designed the experiments. A.M. and C.A.M performed the experiments. A.M. and C.A.M. processed the networks, geneset collections, and analyzed the data. A.M., C.A.M., and A.K. wrote the manuscript. All authors read, edited, and approved the final manuscript.

## Funding

This work was primarily supported by the US National Institutes of Health (NIH) grant R35 GM128765 to A.K.

## Availability of data and materials

The data used in this study, including the gene list-collections and networks, are freely available on GitHub repository *ModGenePlexus*, which additionally contains code to reproduce the results in this study as well as add new gene classification methods is available at https://github.com/krishnanlab/ModGenePlexus. *Conflict of Interest*: none declared.

## Acknowledgements

We thank members of the Krishnan Lab for valuable discussions and feedback on the manuscript.

## Notes

### Competing Interest Statement

The authors have declared no competing interest.

