## Supplemental Material for "A module-based approach for post-omics, post-GWAS network-based gene classification"

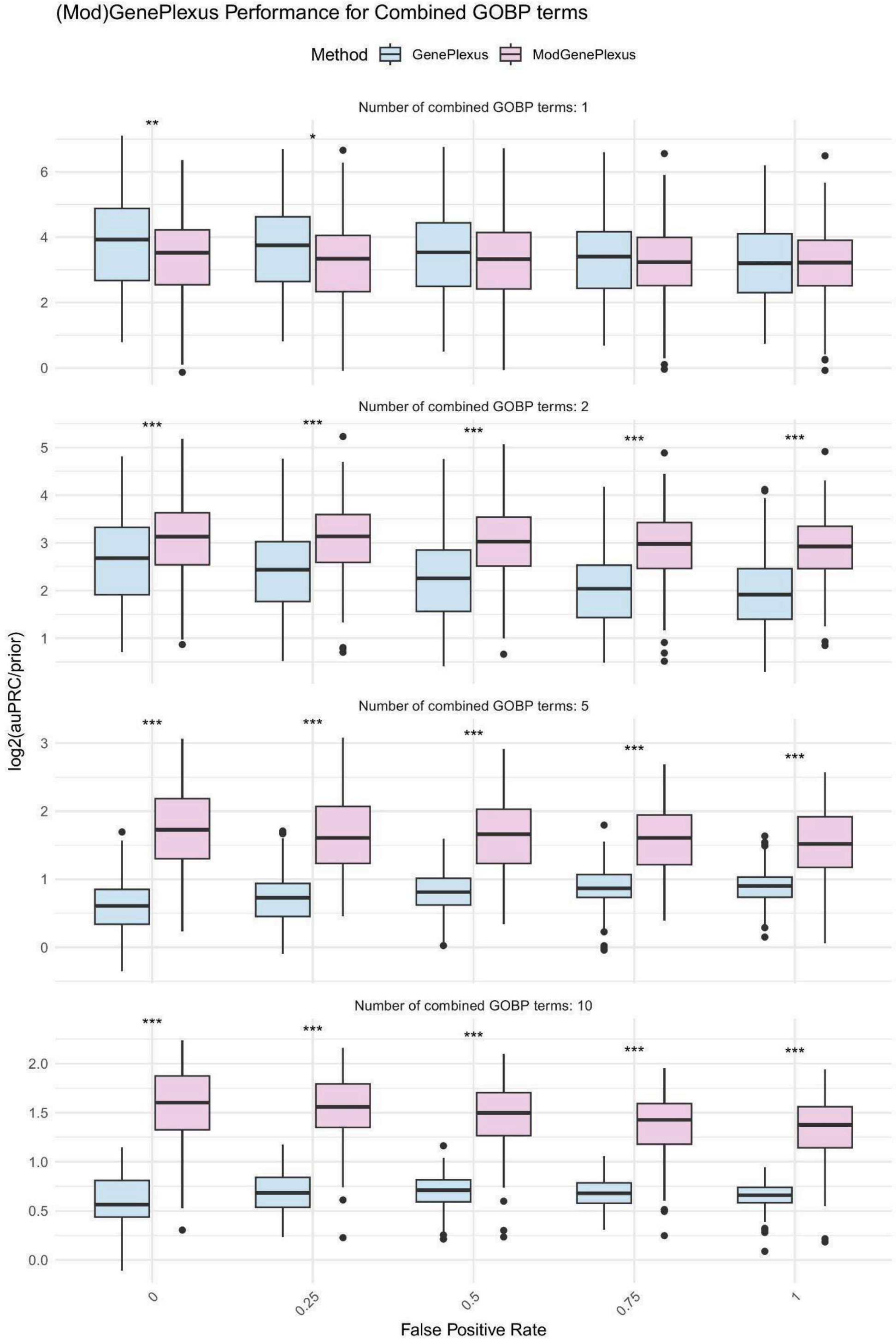


**Figure S1: *ModGenePlexus* outperforms *GenePlexus* on simulated traits generated by combining multiple GOBP terms.** *Performance on simulated traits generated by combining*

*multiple GOBP terms. Models were tested on individual GOBP terms and simulated traits composed of 2, 5, and 10 combined terms. False positives were introduced at rates of 0, 0.25, 0.50, 0.75, and 1.0, corresponding to proportions of the original set size. Results show that as the number of combined GOBP terms increases, the performance gap between ModGenePlexus and GenePlexus widens, with ModGenePlexus consistently outperforming GenePlexus, particularly for multi-term traits.*

**Evaluating additional model steps between *GenePlexus* and *ModGenePlexus***

*ModGenePlexus* introduces several key differences compared to *GenePlexus*. with the most notable being the incorporation of a filtering step during module discovery. This step removes genes that do not belong to sufficiently large modules, ensuring that only robust modules are used for training logistic regression models. Additionally *GenePlexus* is used on top of DOMINO-derived modules. This module discovery process is itself a form of gene classification as genes are propagated that were not part of the initial geneset. Lastly, *ModGenePlexus* creates multiple models – one for each module – rather than a single model for all module genes. We evaluated our method in multiple steps of this pipeline for the transcriptomic datasets. First we run *ModGenePlexus* on modules without their propagated genes, except the propagated positive test genes are included. By including these we tested whether *GenePlexus* is able to distinguish between positive and negative labels better by using the neutral propagated genes, or if *GenePlexus* is only performing better because the semi-supervised learning in DOMINO successfully recovered some positive test genes. We demonstrate that using all propagated genes from DOMINO gives superior results for *ModGenePlexus* (**Figure S2)**. We next assessed a baseline where a single model is created for all module genes – those that are part of the initial disease, in big enough modules, and the propagated genes – rather than creating an individual model for each module. Our results indicate that *ModGenePlexus* significantly outperforms this approach when individual models are created for each module (**Figure S3**).

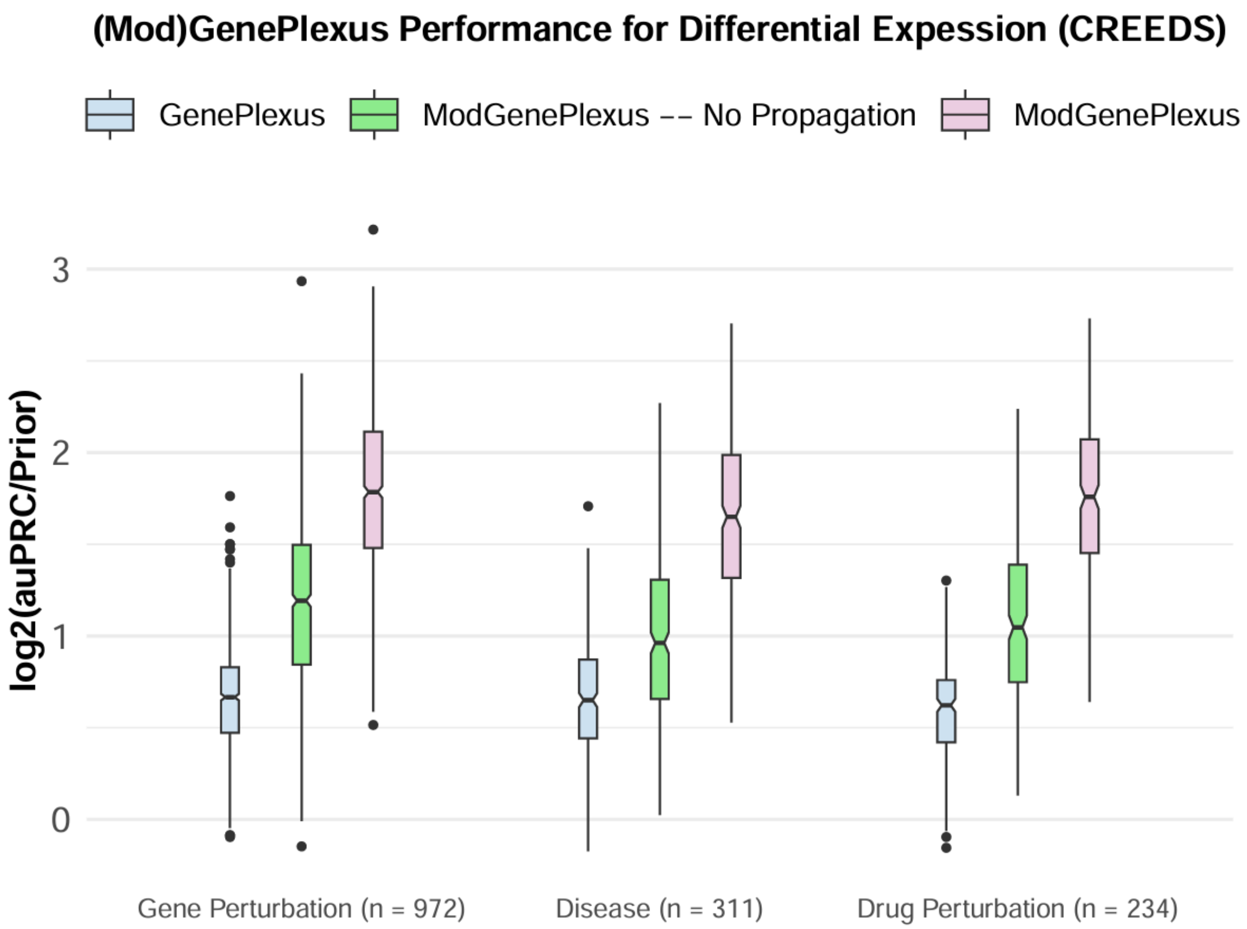


**Figure S2: Full *ModGenePlexus* outperforms *ModGenePlexus* without propagation.** *These plots compare the performance of GenePlexus (blue), ModGenePlexus (pink), and ModGenePlexus without propagation (green). ModGenePlexus performance when not including genes propagated from DOMINO clustering performs worse than the full ModGenePlexus method.*

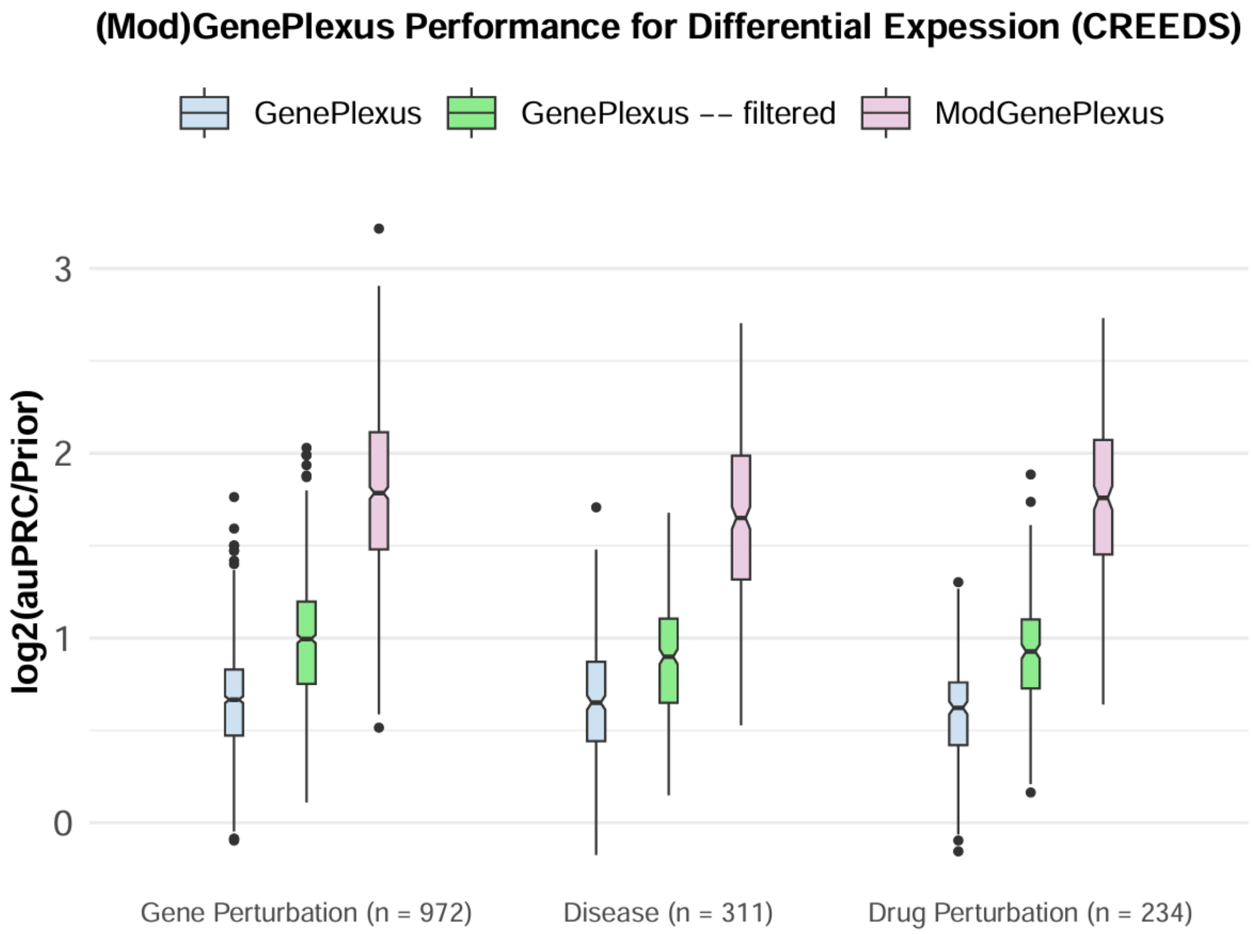


**Figure S3: Full *ModGenePlexus* outperforms *GenePlexus* on an individual list of all module genes.** *GenePlexus performance when only using genes that were assigned to a large enough module (>= 10 genes) in DOMINO is inferior to ModGenePlexus*

**Evaluating Gene Classification with DOMINO vs *ModGenePlexus***
The DOMINO clustering algorithm performs a form of gene classification by propagating genes that are strongly connected to discovered modules. This raises an important question: do *GenePlexus* and *ModGenePlexus* provide added utility beyond what DOMINO alone contributes? To address this, we compared the performance of DOMINO-based gene classification to *ModGenePlexus* by calculating F1 scores using two complementary approaches. First, we computed the F1 score at the maximum point on the auPRC curve **(Figure S4A)**, providing an optimistic view of model performance. Second, we computed F1 scores using the top N predictions from *ModGenePlexus*, where N equals the number of genes propagated by DOMINO after clustering **(Figure S4B)**, providing a stricter and more conservative evaluation. Two evaluations are also useful because it is user-decision what genes to use from *(Mod)GenePlexus* results. To calculate DOMINO's F1 scores, we labeled the propagated test genes – both positive and negative – and calculated precision and recall accordingly. Across both evaluation strategies, *ModGenePlexus* consistently outperformed DOMINO for the majority of transcriptomics disease gene sets. The performance gap was

especially pronounced when evaluating F1 at the auPRC maximum, but even under the more conservative top-N condition, *ModGenePlexus* still demonstrated superior classification accuracy.

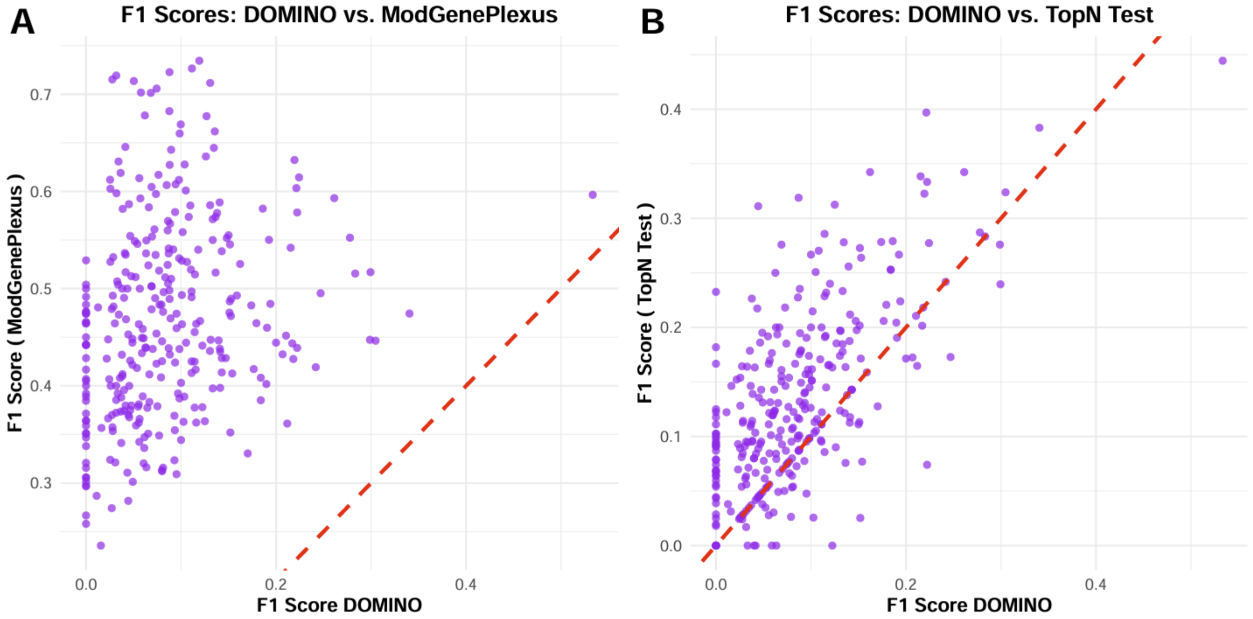


**Figure S4: Comparing F1 scores of DOMINO and *ModGenePlexus* for transcriptomics disease gene sets.** *(A) F1 scores are calculated at the optimal point on the auPRC curve, representing the best-case performance of ModGenePlexus. (B) A more conservative evaluation is shown, where the number of predicted genes from ModGenePlexus is matched to the number of genes propagated by DOMINO. In both settings, ModGenePlexus consistently outperforms DOMINO-based gene classification.*

**Limitations of DOMINO Gene Classification**

A key advantage of *ModGenePlexus* is that it provides predictions for all genes in the network, allowing users to assess genome-wide relationships to their input gene list. In contrast, DOMINO is a binary classification algorithm, meaning it only outputs propagated genes while discarding those removed during clustering. This limits its ability to offer a comprehensive gene classification framework. **Figure S5A** illustrates that for the vast majority of transcriptomics disease datasets, fewer than 10% of positive test genes are recovered during DOMINO's propagation step. Additionally, **Figure S5B** shows that when examining the composition of propagated genes, negative test genes outnumber positive test genes in most cases. This suggests that DOMINO not only fails to recover many disease-associated genes but also propagates a substantial number of negative test genes, reducing classification precision. Furthermore, its lack of genome-wide predictions prevents users from fully contextualizing their input gene sets. These limitations highlight a major advantage of *(Mod)GenePlexus*, which retains genome-wide interpretability while improving classification accuracy.

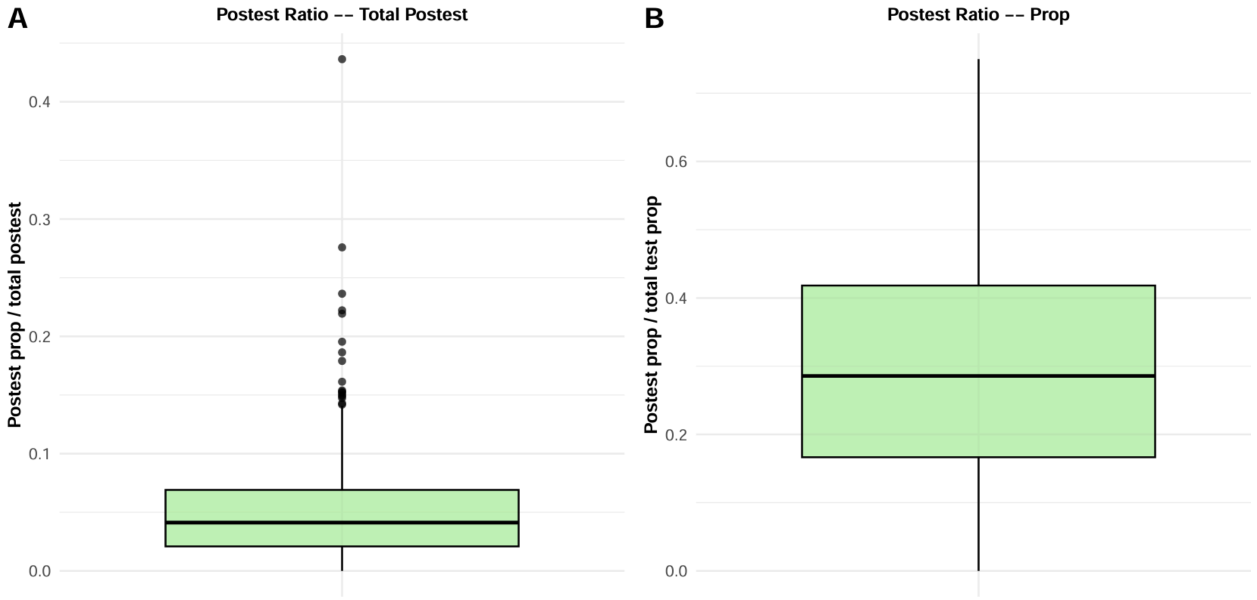


**Figure S5: Domino gene classification alone fails to uncover most known positive genes.** *(A) Proportion of positive test genes recovered by DOMINO relative to the total number of positive test genes in transcriptomics disease datasets. (B) Proportion of positive test genes among all test genes propagated by DOMINO, showing that negative test genes often outnumber positives.*

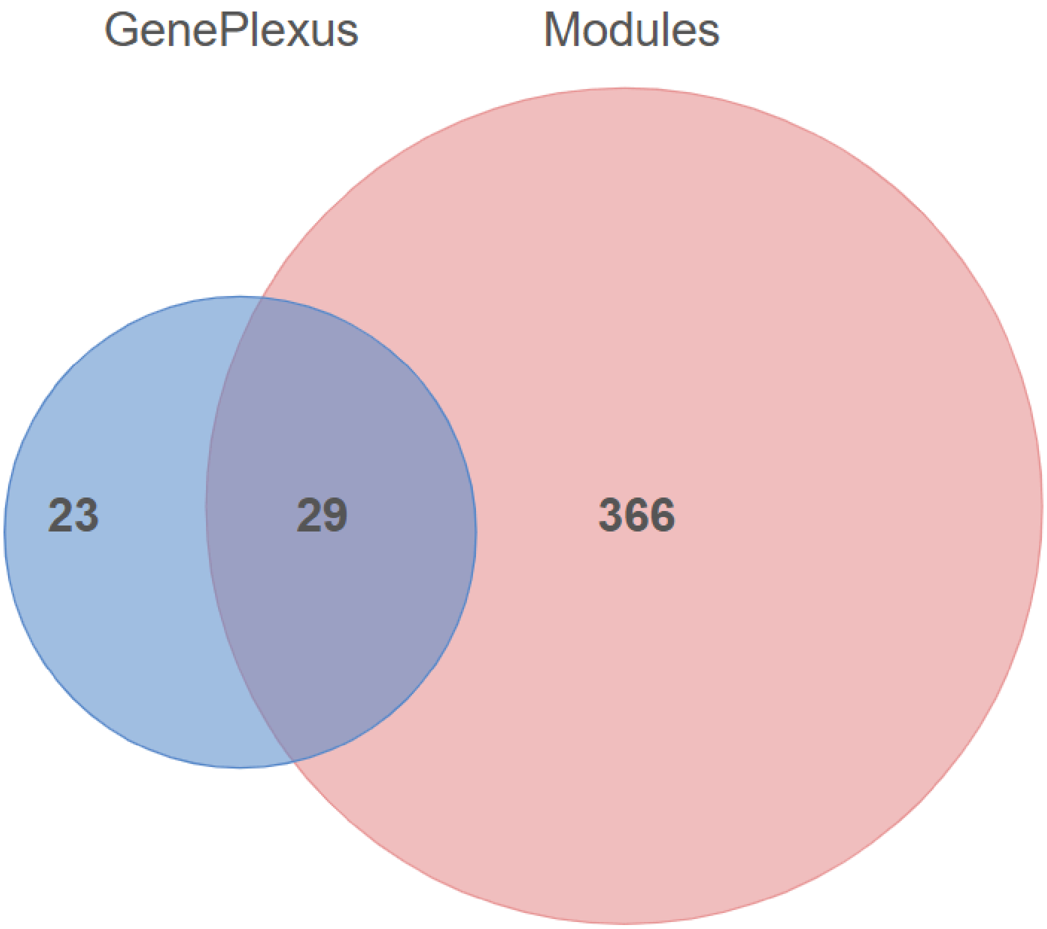


**Figure S6: The number of enriched GOBP terms in *GenePlexus* and modules.** *Enrichment on modules uncovers many more terms than running enrichment on the full initial gene list.*

| GOBPs enriched for Module 1 and 3 | | | | | | | |
|---|---|---|---|---|---|---|---|
| Rank | Description | qvalue | Count | TermSize | Term | GenePlexusRank | GPQvalues |
| 1 | oxidative phosphorylation | 1.02e-170 | 105 | 141 | Module_1 | | |
| 2 | aerobic respiration | 4.20e-168 | 113 | 188 | Module_1 | | |
| 3 | proton motive force-driven ATP synthesis | 2.53e-127 | 66 | 75 | Module_3 | | |
| 4 | mitochondrial respiratory chain complex assembly | 5.64e-110 | 71 | 101 | Module_1 | | |
| 5 | ATP metabolic process | 1.93e-107 | 69 | 131 | Module_3 | 5 | 2.31e-17 |
| 6 | proton transmembrane transport | 1.24e-90 | 64 | 153 | Module_3 | 26 | 3.85e-05 |
| 7 | ribose phosphate biosynthetic process | 3.83e-85 | 68 | 225 | Module_3 | 6 | 2.65e-16 |
| 8 | mitochondrial membrane organization | 8.37e-29 | 30 | 113 | Module_1 | 42 | 3.79e-04 |
| 9 | establishment of protein localization to mitochondrion | 1.49e-23 | 27 | 122 | Module_1 | | |
| 10 | mitochondrial transport | 1.86e-23 | 31 | 182 | Module_1 | 52 | 8.12e-04 |
| 11 | establishment of protein localization to mitochondrial membrane | 8.04e-23 | 18 | 37 | Module_1 | | |
| 12 | protein insertion into membrane | 9.13e-18 | 19 | 75 | Module_1 | | |

**Table S1:** GOBP terms enriched by both module 1 and module 3 for Type 2 Diabetes.

| Top Enriched GOBPs for Module 5 | | | | | | |
|---|---|---|---|---|---|---|
| Rank | Description | qvalue | Count | TermSize | GenePlexusRank | GPQvalues |
| 1 | aerobic respiration | 7.65e-88 | 66 | 188 | | |
| 2 | oxidative phosphorylation | 9.04e-86 | 60 | 141 | | |
| 3 | mitochondrial membrane organization | 6.67e-56 | 42 | 113 | 42 | 3.79e-04 |
| 4 | proton motive force-driven mitochondrial ATP synthesis | 2.16e-55 | 36 | 66 | | |
| 5 | mitochondrial transport | 2.08e-54 | 47 | 182 | 52 | 8.12e-04 |
| 6 | protein localization to mitochondrion | 1.50e-51 | 41 | 128 | | |
| 7 | ATP metabolic process | 2.58e-49 | 40 | 131 | 5 | 2.31e-17 |
| 8 | protein targeting to mitochondrion | 2.30e-48 | 37 | 104 | | |
| 9 | mitochondrial respiratory chain complex assembly | 4.72e-47 | 36 | 101 | | |
| 10 | ribose phosphate biosynthetic process | 1.44e-40 | 41 | 225 | 6 | 2.65e-16 |
| 11 | mitochondrial transmembrane transport | 3.01e-29 | 22 | 58 | 50 | 7.32e-04 |
| 12 | protein transmembrane import into intracellular organelle | 9.87e-29 | 19 | 36 | | |
| 13 | proton transmembrane transport | 1.91e-27 | 28 | 153 | 26 | 3.85e-05 |
| 14 | protein insertion into membrane | 2.11e-26 | 22 | 75 | | |
| 15 | intracellular copper ion homeostasis | 7.77e-12 | 8 | 16 | | |
| 16 | copper ion homeostasis | 4.33e-11 | 8 | 19 | | |
| 17 | copper ion transport | 1.32e-07 | 6 | 18 | | |
| 18 | apoptotic mitochondrial changes | 5.63e-07 | 10 | 104 | | |
| 19 | mitochondrial outer membrane permeabilization | 1.16e-05 | 6 | 36 | | |
| 20 | positive regulation of membrane permeability | 4.69e-05 | 6 | 46 | | |

**Table S2**: The top 20 terms enriched for module 5 for Type 2 Diabetes. This includes the relevant copper ion transport pathway – enriched for module 5 but not for *GenePlexus*.

| Top Enriched GOBPs for Module 1 | | | | | | |
|---|---|---|---|---|---|---|
| Rank | Description | qvalue | Count | TermSize | GenePlexusRank | GPQvalues |
| 1 | oxidative phosphorylation | 1.02e-170 | 105 | 141 | | |
| 2 | aerobic respiration | 4.20e-168 | 113 | 188 | | |
| 3 | proton motive force-driven ATP synthesis | 4.57e-123 | 69 | 75 | | |
| 4 | mitochondrial respiratory chain complex assembly | 5.64e-110 | 71 | 101 | | |
| 5 | ATP metabolic process | 7.21e-100 | 72 | 131 | 5 | 2.31e-17 |
| 6 | ribose phosphate biosynthetic process | 3.95e-75 | 70 | 225 | 6 | 2.65e-16 |
| 7 | proton transmembrane transport | 2.24e-52 | 49 | 153 | 26 | 3.85e-05 |
| 8 | mitochondrial membrane organization | 8.37e-29 | 30 | 113 | 42 | 3.79e-04 |
| 9 | mitochondrial calcium ion transmembrane transport | 2.22e-24 | 15 | 18 | | |
| 10 | establishment of protein localization to mitochondrion | 1.49e-23 | 27 | 122 | | |
| 11 | mitochondrial transport | 1.86e-23 | 31 | 182 | 52 | 8.12e-04 |
| 12 | establishment of protein localization to mitochondrial membrane | 8.04e-23 | 18 | 37 | | |
| 13 | mitochondrial transmembrane transport | 4.01e-20 | 19 | 58 | 50 | 7.32e-04 |
| 14 | protein insertion into membrane | 9.13e-18 | 19 | 75 | | |
| 15 | mitochondrial calcium ion homeostasis | 1.09e-16 | 13 | 26 | | |
| 16 | protein transmembrane import into intracellular organelle | 8.32e-10 | 10 | 36 | | |
| 17 | mitochondrial protein processing | 1.89e-07 | 6 | 13 | | |
| 18 | iron-sulfur cluster assembly | 2.42e-06 | 7 | 30 | | |
| 19 | metallo-sulfur cluster assembly | 2.42e-06 | 7 | 30 | | |
| 20 | regulation of generation of precursor metabolites and energy | 3.28e-05 | 11 | 130 | | |

**Table S3:** The top 20 terms enriched by module 1 for Type 2 Diabetes. This includes the relevant iron-sulfer module assembly pathway – enriched for module 1 but not for *GenePlexus*.

| Iron pathways discovered in Modules | | | | | |
|---|---|---|---|---|---|
| Rank | Description | qvalue | Count | TermSize | Term |
| 1 | iron-sulfur cluster assembly | 2.42e-06 | 7 | 30 | Module_1 |
| 2 | intracellular iron ion homeostasis | 3.74e-04 | 6 | 68 | Module_3 |

**Table S4**: Iron-related pathways found enriched in modules 1 and 3 for Type 2 Diabetes.

| Insulin pathways discovered in Modules | | | | | |
|---|---|---|---|---|---|
| Rank | Description | qvalue | Count | TermSize | Term |
| 1 | response to insulin | 1.83e-07 | 14 | 254 | Module_7 |
| 2 | cellular response to insulin stimulus | 4.10e-06 | 11 | 193 | Module_7 |
| 3 | regulation of insulin-like growth factor receptor signaling pathway | 1.46e-04 | 4 | 23 | Module_12 |

**Table S5:** Insulin-related terms enriched in modules 7 and 12. These were enriched for multiple modules but not for *GenePlexus*.

| Top Enriched GOBPs Using GenePlexus | | | | |
|---|---|---|---|---|
| Rank | Description | qvalue | Count | TermSize |
| 1 | generation of precursor metabolites and energy | 2.51e-33 | 297 | 492 |
| 2 | cytoplasmic translation | 5.48e-28 | 118 | 149 |
| 3 | mitochondrial ATP synthesis coupled electron transport | 1.36e-21 | 80 | 96 |
| 4 | purine nucleoside triphosphate metabolic process | 4.35e-20 | 113 | 160 |
| 5 | ATP metabolic process | 2.31e-17 | 94 | 131 |
| 6 | ribose phosphate biosynthetic process | 2.65e-16 | 138 | 225 |
| 7 | protein folding | 1.85e-11 | 119 | 206 |
| 8 | glucose metabolic process | 1.07e-09 | 107 | 188 |
| 9 | small molecule catabolic process | 9.66e-09 | 180 | 368 |
| 10 | pyruvate metabolic process | 2.35e-08 | 70 | 113 |
| 11 | response to topologically incorrect protein | 4.97e-08 | 88 | 154 |
| 12 | carbohydrate catabolic process | 3.12e-07 | 87 | 156 |
| 13 | establishment of protein localization to membrane | 5.73e-07 | 131 | 262 |
| 14 | protein targeting | 1.73e-06 | 152 | 318 |
| 15 | striated muscle cell development | 1.77e-06 | 47 | 72 |
| 16 | myofibril assembly | 1.80e-06 | 46 | 70 |
| 17 | hexose biosynthetic process | 3.35e-06 | 55 | 90 |
| 18 | fatty acid metabolic process | 5.70e-06 | 174 | 379 |
| 19 | organic acid catabolic process | 8.14e-06 | 121 | 247 |
| 20 | carboxylic acid catabolic process | 8.14e-06 | 121 | 247 |

**Table S6:** The top 20 GOBPs enriched by *GenePlexus* for Type 2 Diabetes.
